# Gut bacterial sphingolipid production modulates dysregulated skin lipid homeostasis

**DOI:** 10.1101/2024.12.29.629238

**Authors:** Min-Ting Lee, Xiaoqing Tan, Henry H. Le, Kevin Besler, Sharon Thompson, Tamia Harris-Tryon, Elizabeth L. Johnson

## Abstract

Sphingolipids are an essential lipid component of the skin barrier with alterations in skin sphingolipid composition associated with multiple skin disorders including psoriasis, atopic dermatitis, and ichthyosis. Contributions to skin sphingolipid abundance are not well characterized, thus the main method of modulating skin lipid levels is the topical application of creams rich with sphingolipids at the skin surface. Evidence that diet and gut microbiome function can alter skin biology proposes an intriguing potential for the modulation of skin lipid homeostasis through gut microbial metabolism, but potential mechanisms of action are not well understood. Sphingolipid synthesis by prominent gut microbes has been shown to affect intestinal, hepatic and immune functions with the potential for sphingolipid-producing bacteria to affect skin biology through altering skin sphingolipid levels. To address this question, we used bioorthogonal chemistry to label lipids from the sphingolipid-producing bacteria *Bacteroides thetaiotaomicron* and trace these lipids to the skin epidermis. Exposing mice to *B. thetaiotaomicron* strains mutant in the ability to produce sphingolipids resulted in significantly lower transfer of gut microbiome-derived lipids to the skin, while also altering skin biology and altering expression of skin barrier genes. Measurement of skin ceramide levels, a class of sphingolipids involved in skin barrier function, determined that skin sphingolipid levels were altered in the presence of gut sphingolipid-producing bacteria. Together this work demonstrates that gut bacterial lipids can transfer to the skin and provides a compelling avenue for modulating sphingolipid-dominant compartments of the skin using sphingolipid-producing bacteria of the gut microbiome.

## INTRODUCTION

The skin is a vast barrier organ that prevents dehydration, functions in thermoregulation, and acts as a physical and immunological barrier against the invasion of pathogens and foreign substances^1,2^. The outermost enucleated layer of the epidermis, termed the stratum corneum (SC), makes critical contributions to these essential functions^3^. In addition to its function in thermoregulation and water balance, the skin also provides an ecological niche for bacteria, fungi, viruses and mites that have evolved to thrive in the desiccant acidic skin landscape and metabolize the lipid and amino acid nutrients generated by the epidermis^4^. These resident microbes are intertwined within the SC^5-7^ with several species shown to promote skin barrier function and repair^8,9^. Lipids are a critical component of the SC, of which ceramides (CERs) are dominant. CERs are a type of sphingolipid which have important structural and signaling functions in skin biology^10^ and host-microbe interactions^11^. Indeed, decreased levels of epidermal CERs are strongly associated with skin disorders such as aging^12,13^, atopic dermatitis^14,15^, ichthyosis^16,17^ and psoriasis^18^. Contributions to the lipid environment of the skin can have multiple functional consequences, however; the comprehensive factors that contribute to the skin lipid environment remain unclear.

Diet is a potentially potent modifier of the lipid composition and functions of the skin. Mice fed a high-fat diet display shifts in skin lipid content^19^, certain diet-induced metabolic disorders have been associated with compromised skin barrier functions^20,21^, and a western diet composed of high sucrose and saturated fat has been found to predispose individuals to inflammatory skin disorders^22^. Taken together, these findings suggest that dietary components may be key regulators of skin lipid composition, barrier function, and skin health. Despite the potential to modify skin biology through diet, relatively little is known about the consequences of diet modification to skin sphingolipid concentrations and the effects these changes may have on the regulation of skin barrier function. Moreover, the microbial community of the gut has the potential to mediate diet-induced changes to skin biology, but more insight is needed regarding the mechanisms governing how the gut microbiome may affect the lipid environment of the skin.

When considering the influence of the gut microbiome on skin sphingolipids, there is evidence that microbial sphingolipids may contribute to skin biology. Fukami and colleagues (2010)^23^, prepared a stable isotope-labeled bacterial dihydroceramide, a CER precursor, by culturing the environmental microbe *Acetobacter malorum* with ^13^C-labeled acetic acid and administering the purified products to mice for 12 consecutive days. They found that the skin showed detectable labeled dihydroceramide-derivatives, demonstrating the transfer of these isolated sphingolipid products from the gut to the epidermis. The epidermis demonstrated the highest intensity signal from the introduced lipids among the surveyed tissues, indicating that an isolated environmental microbe-derived CER precursor can be incorporated into the skin. However, it is still not known if live bacteria in the gut are able to contribute to the sphingolipid environment of the skin, resulting in similar results as observed in the above-mentioned feeding experiment. In addition to effects of diet on skin microbial communities, an expanding research area is starting to focus on the modulating potential of the gut microbiome on diet-dependent host phenotypes in organs distal to the intestinal tract^24-26^.

To determine whether gut microbiome sphingolipid synthesis can influence host lipid metabolism in the skin, we take advantage of our recently developed techniques to trace lipid transfer from the gut microbiome. In this system, we use a bioorthogonal chemistry-based labeling strategy to characterize and trace sphingolipids from the gut commensal *Bacteroides thetaiotaomicron* to the tissues of mice fed a lipid-depleted diet where we have previously observed direct transfer of labeled *B. thetaiotaomicron*-lipids to the host colon^25,27^ and liver^25^. Given that *B. thetaiotaomicron* is one of the major producers of bacterial sphingolipids in the gut, and the importance of CERs/sphingolipids to host skin biology, we hypothesized that gut microbial sphingolipid synthesis can affect host skin phenotypes. To investigate the potential role of gut bacterial sphingolipid production on host skin, we analyzed lipid transfer, the skin transcriptome, and skin sphingolipid concentrations of mice fed a lipid-free diet that were exposed to sphingolipid-competent (BTWT) as compared to sphingolipid-deficient (SLMUT) gut microbes. Exposure to sphingolipid competent microbes resulted in microbial lipid transfer to the skin and alterations in skin ceramide levels that restored the skin biology to a state similar to that before the dietary intervention.

## MATERIALS AND METHODS

### Bacterial strains and culture conditions

*Bacteroides thetaiotaomicron* VPI-5482 (wild-type, BTWT) and the corresponding sphingolipid-null strain (SLMUT) were obtained and prepared as previously described^25,27,28^. In brief, SLMUT is a mutant strain of BTWT with a transposon inserted in a position 88% from the start of gene BT_0870, a homologous gene to a gene with serine palmitoyl transferase (SPT) activity in *Bacteroides fragilis*^*29*^, which renders an inhibition of the canonical sphingolipid synthesis pathway in SLMUT.

Brain heart infusion medium (BHIS) and minimal media (MM) were used to culture BTWT and SLMUT. MM consists of 13.6g KH_2_PO_4_, 0.875g NaCl, 1.125g (NH_4_)2SO_4_, 1 mL hemin solution (500mg dissolved in 10 mL of 1M NaOH, followed by a dilution with water to result in 500 mL final volume), 5g glucose, 1 mL MgCl_2_ (0.1M in water), 1 mL FeSO_4_·7H_2_O, 1 mL vitamin K3 (1 mg/mL in absolute ethanol), 1 mL CaCl_2_ (0.8% w/v), 250 mL vitamin B12 solution (0.02 mg/mL), and 5g L-cysteine hydrochloride anhydrous. pH was adjusted to 7.2 using 5M NaOH. Media were prepared freshly and pre-reduced in an anaerobic chamber (70% N_2_, 25% CO_2_, and 5% H_2_). The inoculation stock was prepared by inoculating either BTWT or SLMUT into pre-warmed (37°C) BHIS and incubated at 37°C under anaerobic condition.

### Bacterial gavage material preparation

The detailed procedure has been published in Le *et al* (2022)^25^. In brief, 1 mL inoculation stock of BTWT and SLMUT was inoculated into 25 mL pre-warmed (37°C) MM supplemented with alkyne tagged sphingolipid precursor, alkyne palmitic acid (PAA), to encourage the assimilation and metabolism of PAA by BTWT/SLMUT to produce sphingolipids/other background lipids (BTWT^PAA^ and SLMUT^PAA^). After overnight incubation at 37°C anaerobically, bacterial pellets were prepared by centrifugation at 4,000 x g for 20 minutes at room temperature, followed by three times washing and pelleting with sterile 1X phosphate-buffered saline (PBS) to remove the residual PAA. PBS was then used to dilute both BTWT^PAA^ and SLMUT^PAA^ to about 10^8^ CFU per 200 µL PBS.

### Laboratory mice and diet

For the transcriptomic experiment, 32 mice contributing to the experiment were allocated evenly to each treatment group with 4 mice per cage and 2 cages per group. For the metabolomics and phenotyping experiments, 24 mice were contributed, with 3 mice per cage and 2 cages per group.

All the experimental mice were received from Taconic Biosciences as conventionally raised, excluded flora, five-week-old, female Swiss Webster mice. Mice were acclimated for five days after delivery, and then weighted and randomly assigned to 4 treatment groups. Mice were kept in autoclaved microisolator cages and housed in humidity and temperature-controlled rooms on a 12-hour light-dark cycle. Autoclaved sterile drinking water was provided *ad libitum* to all the mice, and 2 cages of mice (CHOW group) were fed a breeder chow diet (5021 LabDiet) containing 23.7%-kcal fat, 53.2% carbohydrate, 23.1% protein; the other 6 cages of mice (Vehicle, BTWT^PAA^, and SLMUT^PAA^) received a lipid-free diet (Envigo, TD. 03314) containing 0%-kcal fat, 75.9% carbohydrate, and 24.1% protein for 4 weeks before oral administration of *B. thetaiotaomicron* strains. On the fifth week, two groups of mice were subjected to daily oral administration of either BTWT^PAA^ or SLMUT^PAA^, with the Vehicle group receiving a PBS oral gavage as a control, and the CHOW group being scruffed for ∼30 seconds (equal time per mouse gavage time) to remove any potential bias effects derived from scruffing.

### Skin histology

After shaving the hair, mice dorsal skin tissues were excised and placed between two pieces of 2 cm^2^ filter paper (Whatman) that were pre-soaked in 10% buffered-neutral formalin (pH 7.2) followed by a 48 h fixation at 4°C. The fixed tissue slices were embedded horizontally in the biopsy cassette (VWR) and submitted to the Histology Laboratory of the Animal Health Diagnostic Center at Cornell University for subsequent paraffin embedment, section and hematoxylin and eosin (H&E) staining. Histopathological phenotypes were examined by light microscopy (Leica DM500. Leica, Buffalo Grove, IL), and images were taken at 5X magnification.

### Click chemistry-based detection of microbially-derived lipids in the skin

Skin was collected from the dorsal region and incubated in OCT compound on ice before moved to a -80°C freezer to fully embed tissue in preparation for cryosectioning. Skin was cross sectioned using a cryostat (Leica). DNA was stained with 1 μg/mL Hoechst 33342. *Bacteroides*-derived lipids were detected with AlexaFluor 647 azide (ThermoFisher) using a Click & Go kit (Vector Laboratories) to link the fluorophore to any alkyne-containing metabolites through the click reaction. Images were acquired using fluorescence microscopy (Leica DM500. Leica, Buffalo Grove, IL) at 40X magnification.

### Skin sphingolipid profiling

After shaving the hair, mice skin was collected from the dorsal region and the subcutaneous fat was trimmed off using a scalpel. The rest of the skin was floated on the 1X PBS prepared dispase (1mg/mL) for 2h at 37°C to separate epidermis from the dermis. The epidermal layer was collected by lifting the sheet using autoclaved forceps and cut into small pieces in 2 mL screw-top tube prefilled with 1.0 mm Zirconium beads (OPS Diagnostics). 50 mg samples were then frozen in liquid nitrogen and lyophilized for 2 days. 1 mL HPLC-grade methanol (Fisher Scientific) was added to the dried samples, followed by three rounds of 1 min bead beating in a bead beater homogenizer (BioSpec products). Tubes were inserted into ice in between rounds to prevent lipid deterioration. Samples were further extracted using Qsonica ultrasonic processor with 3s on and 2s off at 100% power for 10 min, followed by overnight methanol extraction on an end-over-end rotor. Samples were then centrifuged at 18,000 x g for 30 minutes at 4°C and the clarified supernatant (∼850 μL) was transferred to a clean 1.5 mL tube. The supernatant was then concentrated to dried metabolome using a SpeedVac vacuum concentrator (Thermo Fisher Scentific), and 200 μL HPLC-grade methanol was added to resuspend the metabolome and subjected to LC-MS/MS analyses.

A targeted skin CERs screening method was established according to Kawana et al. (2020)^30^ and Franco et al. (2018)^31^. Ceramide species in four dominant murine skin ceramide classes, including CER[NDS], a ceramide class consisting of non-hydroxy fatty acids and sphinganines; CER[OS], a ceramide class consisting of ω-hydroxysphingosine; CER[EOS], a ceramide class consisting of ester-linked non-hydroxy fatty acids; and CER[NS], a ceramide class consisting of non-hydroxy fatty acids and sphingosine, were selected to analyze^30^. The m/z values of parent ions (first quadrupole, Q1) and product ions (third quadropole, Q3), and collision energy applied are summarized in Supplemental Table 1. Selective reaction monitoring (SRM) was performed on a Thermo Scientific Vanquish Horizon UHPLC System coupled with a Thermo Scientific TSQ Quantis Triple Quadrupole mass spectrometer equipped with a HESI ion source. Mobile phase A was 78.6% water, 20% acetonitrile, and 0.4% formic acid (v/v). Mobile phase B was 47.8% methanol, 47.8% acetonitrile, 4% chloroform, and 0.4% formic acid. 5 µL of extract was injected and separated on a mobile phase gradient with an Agilent Technologies InfinityLab Poroshell 120 EC-C18 column (50 mm × 2.1 mm, particle size 2.7 μm, part number: 699775-902) maintained at 50°C. Mobile phase A and B gradient started at 10% B for 1 min after injection and increased linearly to 100% B at 9 min and held at 100% B for 10 min, using a flow rate 0.6 mL/min. Absolute intensity was obtained through Xcalibur v4.4.16.14 (Thermo Fisher Scientific) and used to represent and compare the quantity of the detected ceramide species between samples and groups.

### RNAseq library construction and sequencing

After shaving the hair, mice skin was collected from the dorsal region. About 20 mg of the skin tissue was stored in the TRIzol (Invitrogen, Carlsbad, CA) in a RNase-free 2 mL screw-top tube prefilled with 1.0 mm Zirconium beads (OPS Diagnostics) at 4°C for 2 days to ensure the reagent was exposed to the tissue. The skin tissue was homogenized with three rounds of 1 min bursts of bead beating in a bead beater homogenizer (BioSpec products), and tubes were inserted into ice in between rounds to prevent RNA degradation. Chloroform was added to the lysate for phase separation *as per* manufacturer’s instructions. The top layer was further cleaned and concentrated with the RNA Clean & Concentrator Kit (Zymo Research) in combination with the on-column DNase treatment from the kit to remove the potential left-over genomic DNA following the manufacturer’s instructions. After checking the quality and quantity, RNA libraries were prepared using the QuantSeq 3’ mRNA-Seq Library Prep kit FWD for Illumina (Lexogen, Vienna, Austria) following the manufacturer’s instructions. Samples were sequenced on the Illumina NextSeq500 Platform with 75bp single-end read.

### RNAseq data analysis

Raw read data was first quality-checked using FastQC (v 0.11.8). Transcript abundances were quantified using pseudo-aligner Salmon (v1.4.0) against the cDNA of the *Mus musculus* GRCm39 database. The gene-level differential expression was performed by passing Salmon quantification results to DESeq2 analysis using DESeqDataSetFromTximport function in the R package DESeq2^32^. Heatmaps were plotted using the R package ComplexHeatmap^33^ to present the differential expressed genes (DEGs) between groups. Functional enrichment analysis was performed using the R package clusterProfiler^34^. Dot plots of biological process in the Gene Ontology database (GO-BP) were plotted using R function enrichGO in the clusterProfiler package. Gene lineage network plots for the enrichment results of over-representation tests were generated using the R function cnetplot in the enrichplot package^35^.

### Statistical analysis

All statistical analyses were performed using R (v3.5.1) and GraphPad Prism 9.0 software (GraphPad Software, Inc., La Jolla, CA, USA). One-way ANOVA with Tukey’s multiple correction test was used to compare across groups. For differentially abundant taxa, we applied ALDEx2 (v1.12.0)^36^ at the ASV level and focused on an overall relative abundance >1% and excluded low-abundance taxa from the analysis. ALDEx2 performs centered log ratio transformation on the count data for a compositionally coherent inference and estimates *P* values and false discovery rate (FDR) from independent testing of Monte Carlo Dirichlet instances. FDR < 0.05 was set as the threshold to identify differentially abundant taxa. For RNAseq, The differentially expressed genes (DEGs) were calculated using the DESeq2 package in R with the cutoff values of Benjamini-Hochverg adjusted p value (p.adjust) < 0.05 and absolute log_2_ fold change > 1.

## RESULTS

### Gut bacterial sphingolipid synthesis results in increased levels of lipid transfer to the skin and amelioration of observed losses of subcutaneous fat on a lipid-free diet

To discern specific contributions to host biology derived from gut bacterial sphingolipids, we leveraged our ability to metabolically label lipids in bacteria and monitored for lipid transfer after oral introduction of these labeled cultures into mice^25,37^ (Figure 1). In this model, wild-type *Bacteroides thetaiotaomicron* (BTWT) is cultured with a proxy of palmitic acid (palmitic acid alkyne, PAA) (Figure 1A). The alkyne modification produces trackable lipids that have previously allowed us to confirm the production of sphingolipids derived from BTWT treated with PAA (BTWT^PAA^)^25^. As a control, we also culture PAA with a strain of *B. thetaiotaomicron* that cannot generate sphingolipids (SLMUT), this mutant is able to incorporate PAA into phospholipids but is unable to make sphingolipids from PAA^25^. This allows us to compare BTWT^PAA^-treated mice to SLMUT^PAA^-treated mice and draw conclusions regarding the impact of *B. thetaiotaomicron*-derived sphingolipid synthesis on host lipid metabolism. Additionally, mice were fed a lipid-free diet for 28 days before administration of the *B. thetaiotaomicron* strains to reduce the disturbance of dietary lipids on the PAA-based lipid tracing (Figure 1B). *B. thetaiotaomicron* strains labeled with PAA were administered to mice daily by gavage for 7 days. Potential transfer of *B. thetaiotaomicron* lipids to host tissue, skin biology, skin gene expression, and skin lipid levels were measured to determine if gut microbial sphingolipid synthesis can influence sphingolipid-dependent biological processes in the skin.

**Figure 1.**
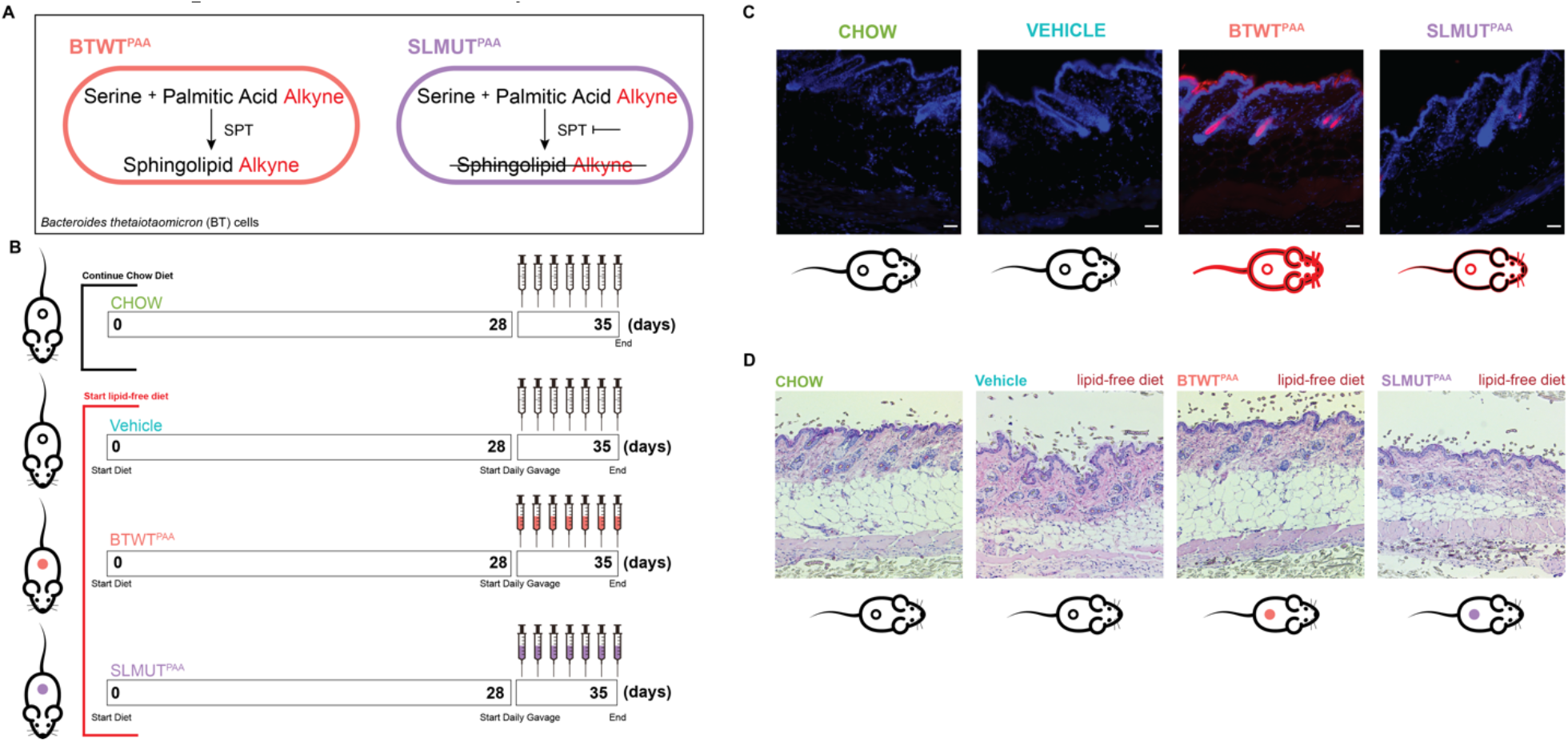
Lipid transfer from the gut microbiome to the skin is dependent on gut microbial sphingolipid synthesis and affects skin morphology including subcutaneous fat. (A) Model of sphingolipid synthesis in BTWT as compared to SLMUT when these *B. thetaiotaomicron* strains are exposed to palmitic acid alkyne (PAA). SPT = serine palmitoyltransferase. (B) Schematic of experiment detailing the diet treatment and *B. thetaiotaomicron* strain gavage over the course of the 35 days from the beginning of the diet intervention to the terminal colletion of samples. (C) Fluorescent microscope images of skin sections with alkyne-bearing microbial lipids detected using AlexaFluor 647 azide (red) and DNA was stained using Hoechst 33342 (blue). Scale bar = 35μm. Mouse icons underneath images have circles that indicate the alkyne-labeling of the gut microbes with the black circles indicating that no alkyne labeled lipids were introduced into the microbiome and red circles indicating that the introduced microbial strains were producing alkyne-labeled lipids. Red outlining of the mouse icon indicates that there was transfer of microbial lipids observed. (D) Microscopy images of H&E stained skin sections. Mouse icons underneath images have circles that indicate the microbial treatment of the mouse with empty cicles indicating no treatment, pink circles indicatitng BTWT^PAA^ treatment, and purple cicles indicating SLMUT^PAA^ treatment.

To test whether gut microbial lipids produced by sphingolipid synthesis-competent microbes are transferred to the sphingolipid-rich stratum corneum region of the skin, we used click chemistry to ligate a fluorophore (AlexaFluor 647 – red) to any alkyne containing lipids derived from BTWT^PAA^. The ligation of the fluorophore occurs post-tissue collection so that the fluorophore does not interfere with the native biochemistry of the system. This click chemistry-based detection of alkyne-containing *B. thetaiotaomicron*-derived lipids demonstrated that gut microbial lipids can be readily transferred to skin (Figure 1C). We observed that *B. thetaiotaomicron*’s capability to produce sphingolipids affects the transfer of *Bacteroides*-derived lipids to the skin as BTWT^PAA^ associated mice demonstrated more detectable gut microbiome-derived skin lipids as compared to SLMUT^PAA^ associated mice (Figure 1C). After oral gavage of PAA-incubated bacterial cultures, no detectable labeled lipids were observed in the stool of mice which suggests that detected alkyne lipids are from the oral introduction of *B. thetaiotaomicron* strains and not the caging environment. To understand if this differential gut microbial lipid transfer has any associations with altered skin biology, we performed H&E staining on the parafilm-sectioned skin of each group of mice. Interestingly, morphological analysis (Figure 1D) revealed that the dermal subcutaneous fat layer is thinner in lipid-free diet fed mice (Vehicle) mice compared to chow-fed (CHOW) mice. Thus, indicating a diet-induced pathology of the skin that can be added to observed hepatic steatosis^25^ incurred on the lipid-free diet. Of note, we also observed that BTWT^PAA^ was the only lipid-free diet-fed group that displayed a subcutaneous fat layer similar to the chow-fed mice (Figure 1D). This suggests that gut microbial sphingolipid synthesis can rescue diet-induced pathologies of the skin.

### *B. thetaiotaomicron* sphingolipid production restored expression of skin barrier regulatory genes and increased certain ceramide levels in the skin of mice fed a lipid-free diet

To further understand potential mechanisms that allow for microbial sphingolipid synthesis to affect skin biology, we used transcriptomics (RNAseq) to examine the genes that were differentially expressed (DEGs) between mice in our diet and *Bacteroides* intervention conditions. Unsupervised clustering using principal component analysis (PCA) was able to distinguish the skin transcriptome of BTWT^PAA^ from other mice fed the lipid-free diet (Vehicle and SLMUT^PAA^) (Figure 2A). Additionally, PCA determined that gene expression in the skin of Chow fed mice and BTWT^PAA^ mice was more similar than mice that were solely put on the lipid-free diet or on the lipid-free diet plus SLMUT^PAA^ (Figure 2A). To determine important gene expression pathways affected by the lipid-free diet and modulated by gut bacterial sphingolipid synthesis, we analyzed DEGs simultaneously using gene ontology (GO) and gene lineage network analysis. Over 300 DEGs (Supplemental Figure 1A) were significantly different between diet conditions for multiple biological functions (Supplemental Figure 1B). The top five functional categories represented were: fatty acid metabolic process, lipid biosynthetic process, monocarboxylic acid metabolic process, acetyl-CoA metabolic process, and acetyl-CoA biosynthetic process (Supplemental Figure 1B, 1C). Therefore, dietary lipid depletion resulted in the downregulation of lipid synthesis and processing genes in the host skin.

**Figure 2.**
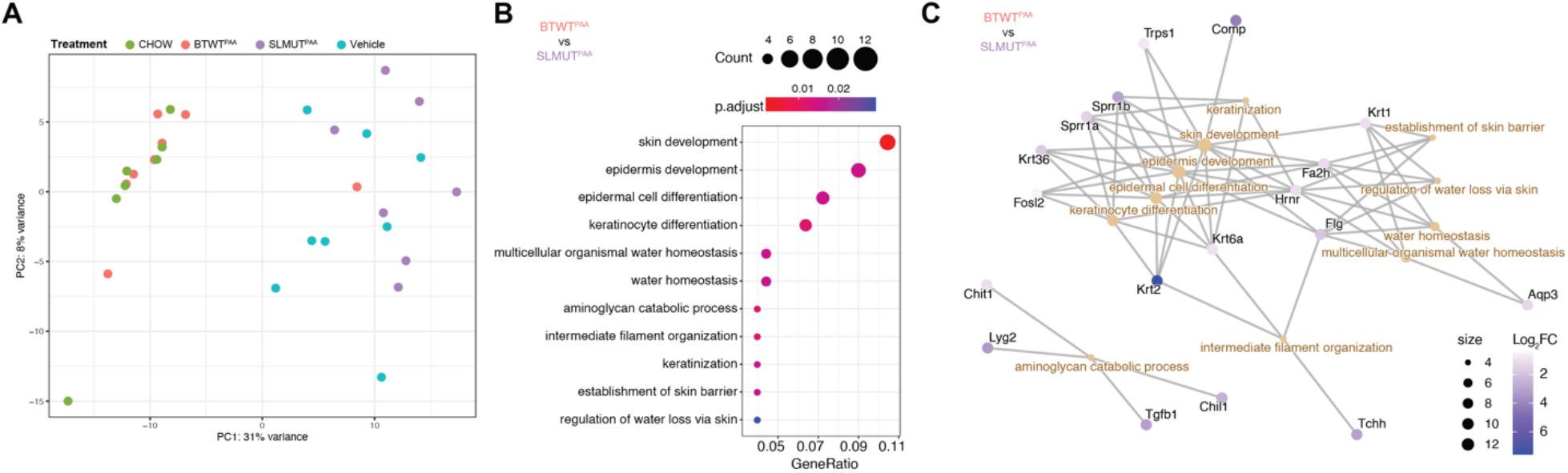
Expression of skin barrier and differentiation genes are enriched in mice supplemented with sphingolipid producing bacteria versus mice supplemented with the isogenic strain that is unable to produce sphingolipids. (A) Principal-component analysis of skin transcriptomes from diet and *Bacteroides*-treatment groups. (B) Biological processes in gene ontology analysis (GO-BP) comparing between skin transcriptomes of BTWT^PAA^ and SLMUT^PAA^ supplemented mice that are significant at q<0.05 ordered by enrichment values, calculated as the portion of genes under selection in the processes over the portion of genes significant in all GO-BPs. Points are color-filled by q-value and sized according to the gene numbers including in the corresponding GO-BP. (C) Gene lineage network plot shows the correlation among DEGs and GO-BP. Points are color-filled by the log_2_ fold change (Log_2_FC) of genes compared between BTWT^PAA^ and SLMUT^PAA^.

To further characterize gene expression programs regulated by *Bacteroides*-derived sphingolipids, we compared transcriptomes between BTWT^PAA^ and SLMUT^PAA^ treated mice and BTWT^PAA^ compared to vehicle treated mice. Multiple genes regulating barrier functions were significantly upregulated in BTWT^PAA^ compared to SLMUT^PAA^ treated mice, including skin development, keratinocyte differentiation, and regulation of trans-epidermal water loss (Figure 2B and 2C). Additionally, genes involved in lipid processing, skin development, and temperature homeostasis were upregulated in BTWT^PAA^ compared to vehicle treated mice (Supplemental Figure 2). These differences in gene expression based on the capacity of the microbiome to produce sphingolipids suggest that gut *Bacteroides* sphingolipid synthesis can influence sphingolipid-related functions of the skin. We also performed gene set enrichment analysis (GSEA) on the DEGs between BTWT^PAA^ and SLMUT^PAA^ using the Hallmark gene sets, a database that represents well-defined biological processes with coherent expression. This analysis demonstrated two gene sets involved in skin repair (wound healing) that were enriched in BTWT^PAA^ mouse skin including transforming growing factor-β (TGF-β) signaling and epithelial mesenchymal transition (EMT)^38^ (Supplemental Figure 3). Consistent with these results, SLMUT^PAA^ displayed suppression of TGF-β expression^39^ and activation of NF-κB, a marker of skin inflammation^40^. In agreement with our findings, one recent study^41^ identified that blocking sphingolipid synthesis in human dermal fibroblasts downregulated genes involved in TGF-β, such as *Acta2* and *Col1a1*, while promoting fibroblast growth factor 2 (*Fgf2*) activation, the antagonistic program to TGF-β (Supplementary Figure 4B-D). Collectively, these results reveal potential mechanisms governing the ability of bacterial sphingolipid production to regulate skin homeostasis.

Focusing on sphingolipid synthesis-specific changes in skin gene expression that may affect skin lipid composition, we extracted BTWT^PAA^/SLMUT^PAA^ DEGs included in the Sphingolipid Metabolism gene set based on the GO database and compared the z-score normalized gene expression across the four groups. Here we centered around the synthesis of sphingolipids in the four main classes dominantly identified in the epidermis of the skin tissue^30^, including non-hydroxy dihydroceramides (CER[NDS]), ω-hydroxydihydroceramides (CER[OS]), esterfied ω-hydroxyceramides (CER[EOS] - also known as acylceramides) and non-hydroxy ceramides (CER[NS]). We identified that the expression of 4 DEGs in the CER[EOS] biosynthetic pathway, including *Aldh3b2, Cers3, Degs1*, and *Pnpla1*, were consistently higher in either BTWT^PAA^ or CHOW-fed conditions as compared to Vehicle-treated and SLMUT^PAA^ conditions (Figure 3A and 3B, green-colored genes). Meanwhile, 4 DEGs involved in fatty-acid processing prior to N-acylation of the sphingoid base were more abundant in Vehicle-treated and SLMUT^PAA^ conditions as compared to BTWT^PAA^ or CHOW-fed conditions (Figure 3A and 3B purple-colored genes).

**Figure 3.**
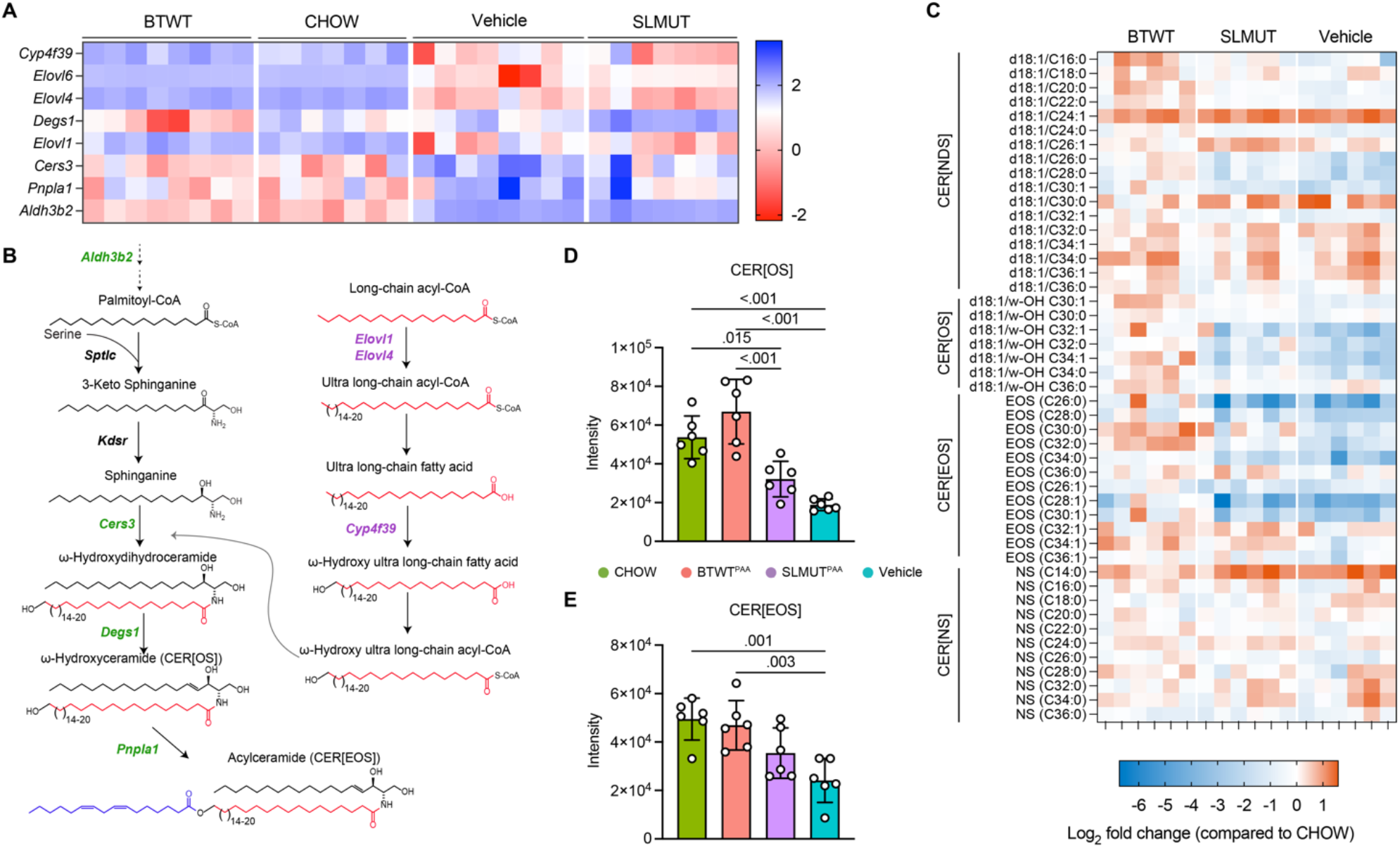
*B. thetaiotaomicron* sphingolipid production in the mouse gut resulted in the enrichment of genes and sphingolipids involved in diverse facets of epithelial barrier function. Mice were fed either chow (CHOW) or a lipid-free diet; and mice fed with the lipid-free diet were orally gavaged with either Vehicle, BTWT^PAA^, or SLMUT^PAA^ (n = 8 in either BTWT^PAA^ or Vehicle; n = 7 in either CHOW or SLMUT^PAA^. (A) Heatmap showing row normalized upregulated and downregulated differentially expressed genes (DEGs), belonging to the Sphingolipid Metabolism gene set, identified between BTWT^PAA^ and SLMUT^PAA^. The heatmap was extended to include all groups. The color key from blue to red indicates low to high gene expression level. Gene names are listed on the left. (B) Scheme of the biosynthesis pathway for ceramides, CER[OS], CER[EOS], and long-chain fatty acids in the epidermis. Genes encoding the corresponding step are annotated on the left-hand side of the arrow. Green-colored genes are upregulated in CHOW and BTWT^PAA^; and purple-colored genes are upregulated in Vehicle and SLMUT^PAA^. (C) Levels of CER[NDS]: ceramide class consisting of non-hydroxy fatty acids and sphinganines, CER[OS]: ceramide class consisting of ω-hydroxysphingosine, CER[EOS]: ceramide class consisting of ester-linked non-hydroxy fatty acids, and CER[NS]: ceramide class consisting of non-hydroxy fatty acids and 4-sphingenines in the mice epidermis. Quantification was performed via LC-MS/MS. The quantities were first normalized to the dry weight of epidermis, and further normalized post-data acquisition by calculating the fold change over the CHOW condition. Values presented as gradient colored heatmap of fold change in log_2_ CONDITION *vs*. CHOW. (D) Total CER[OS] levels in each group, calculating by summing up sphingolipid species identified in this class. (E) Total CER[EOS] levels in each group, calculating by summing up sphingolipid species identified in this class.

### Skin ceramide levels are affected by gut bacterial sphingolipid synthesis

To better understand how gut microbial sphingolipid synthesis may affect observed changes in gene expression and biology through sphingolipid homeostasis, we performed targeted LC-MS analysis of skin lipids. Given the significant difference in expression of genes involved in the regulation of trans-epidermal water loss between BTWT^PAA^ and SLMUT^PAA^ mice (Figure 2D) and that sphingolipids in the SC are regarded as the most critical lipids facilitating this skin barrier function in the epidermis^42^, we postulated that sphingolipid content in the epidermis would differ between BTWT^PAA^ and SLMUT^PAA^ treated mice We measured the abundance of the four main CER classes dominantly identified in the epidermis of the skin tissue^30^, including non-hydroxy dihydroceramides (CER[NDS]), ω-hydroxydihydroceramides (CER[OS]), acylceramides (CER[EOS]) and non-hydroxy ceramides (CER[NS]), using LC-MS/MS. To simultaneously compare between diet conditions and microbial treatment groups, we normalized the sphingolipid content in the skin of the three groups of mice fed the lipid-free diet to mice on the chow diet (CHOW). As shown in Figure 3C, significant global changes to sphingolipid levels from levels observed in the skin of CHOW-fed mice were observed and reflected overall changes in the gene expression levels of sphingolipid synthesis and processing genes.

To match the gene regulation results to the sphingolipid quantification, we summarized all the detectable species of CER[NDS], CER[OS] and CER[EOS], respectively (Figure 3D and 3E, and Supplementary Figure 4). Though no significant difference was found in CER[NDS] among the groups (Supplementary Figure 4), CER[OS] amounts were higher in BTWT^PAA^ compared to the other two groups fed with the lipid-free diet (Figure 3D); and CER[EOS] amounts were consistently lower in Vehicle-treated as compared to either BTWT^PAA^ or CHOW-fed mice (Figure 3E). Collectively, these results suggest that a series of skin barrier-related and sphingolipid synthesis targeted gene expression changes occurred in lipid-free diet fed mice supplemented with BTWT, potentially contributing to increased levels of CER[OS] and CER[EOS].

## DISCUSSION

The SC of mammalian skin is one of the largest sphingolipid reservoirs in the body and serves as a structural and immunological barrier. The gut commensal *Bacteroides* is another key source of sphingolipids to the host and has been found to influence host colon immune development^43,44^ and hepatic lipid metabolism^25,27^ through its de novo production of sphingolipids. Thus, there was potential for microbial influence of skin biology through sphingolipid-dependent mechanisms. Though the skin and gut environments are spatially separated organs of the body, we have demonstrated a mechanistic connection between these two sites through the tracking of gut microbiome-derived lipids to the epidermis. Here, we determined contributions of gut bacterial sphingolipid production to host skin biology, including gene expression, structure, and ceramide homeostasis. We observed that dysregulated skin lipid metabolism and the loss of subcutaneous adipose layer induced by a lipid-free diet were mitigated by supplementing mice with the sphingolipid-producing gut microbe *B. thetaiotaomicron*. Additionally, gut *B. thetaiotaomicron* sphingolipid production regulated genes involved in epidermal water balance, antimicrobial defense, differentiation and development that are tightly connected to the integrity of skin barrier function in mice. Moreover, supplementation with sphingolipid competent *B. thetaiotaomicron* resulted in altered levels of sphingolipid processing genes involved in the production of epidermis prominent sphingolipids, ω-hydroxyceramides and acylceramides. These ceramides can comprise up to 50% of the lipid content of the skin and deficiencies of these complex ceramides are driving factors of skin barrier disruption in skin disorders. The potential to modulate the composition of these skin ceramides using the microbiome opens a new avenue of how to modify the lipid composition of the epidermis to improve skin function.

Studies have shown the potential of supplementing dietary sphingolipids to a variety of mice models in the improvement of skin barrier function^45,46^. For example, Hasegawa *et al* (2011)^47^ administered glucosylceramide (GluCER) orally to hairless mice after 14 days of barrier perturbation induced by ultraviolet B irradiation. They found a significant reduction of transepidermal water loss in the mice treated with GluCER compared to controls, accompanied by increased gene expression of transglutaminase, an enzyme central to the formation of the cornified envelope of the SC. In addition to GluCER, Duan *et al*. (2012)^48^ found that supplementation of sphingomyelin, a type of sphingolipid, upregulated genes encoding ceramide synthases (*Cers*) 3 and 4 in the epidermis of an atopic dermatitis-like skin mouse model and recovered damaged skin barrier functions induced by tape-stripping. In our study, we identified a BTWT-treatment dependent upregulation of not only *Cers3*, but also *Degs1* and *Pnpla1*, which are necessary for the production of CER[OS]^49^ and CER[EOS]^50^ respectively. Importantly, we also identified that this effect co-occurred with the transfer of microbiome derived lipids to the skin. Recently, one study^51^ supplemented human subjects with food with acetic acid bacteria containing dihydroceramide for 12 weeks and showed a significantly improved SC hydration without any harmful effects. Our findings provide potential mechanisms that allow for gut microbes to have similar effects on skin phenotypes that have been observed in the supplementation-based studies mentioned above. Specifically, we showed that sphingolipid production from the gut commensal *B. thetaiotaomicron* contributes to the gene regulation machinery underlying the skin water retention and barrier functions in the lipid-free diet fed mice. Overall, our current work describes new ways that gut commensal metabolic function can affect skin biology. The tracing of bacterial lipids from gut microbes to the skin opens a new avenue to target lipid-dependent skin functions in a mechanistic manner. These insights could be critical in further developing promising microbiome-centric methods of modulating skin biology in the context of skin-based disorders that are modulated by sphingolipid/ceramide composition and abundance in the stratum corneum.

## ACKNOWLEDGEMENTS

Research in the Johnson laboratory is supported by the National Institutes of Health (NIH) through an Early-Stage Investigator Maximizing Investigators’ Research Award (ESI-MIRA) to ELJ (R35GM138281). ELJ is a Pew Biomedical Scholar and a Schwartz Award Visionary Scientist Awardee. Research in the Harris-Tryon lab is supported by the National Institutes of Health grant NIAMS-K08AR076459 and the Burroughs Wellcome Fund 1022777 to TAH. TAH is the Thomas L. Shields Professor of Dermatology at UT Southwestern.

