## Supplementary material for "Gut bacterial sphingolipid production modulates dysregulated skin lipid homeostasis": Supplmental Information

### SUPPLEMENTAL FIGURES & TABLES

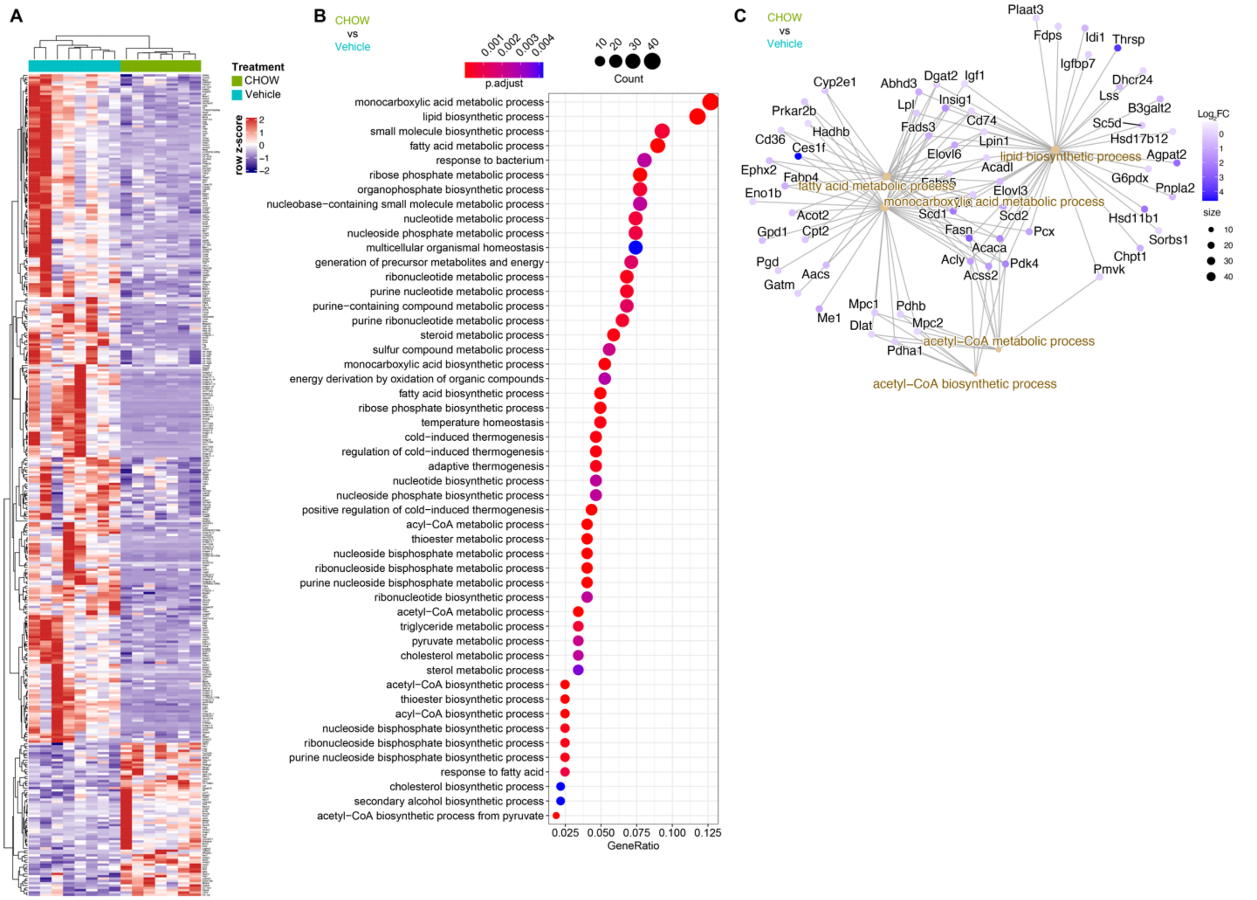


**Supplemental Figure 1.** (A) Heatmap displaying 354 significantly differentially expressed genes in the skin between mice fed the CHOW diet and mice fed the lipid-free diet (Vehicle). (B) The top 50 biological processes from gene ontology analysis (GO-BP) that are significantly differentially expressed between CHOW diet and lipid free-diet (Vehicle) fed mice at an adjusted p-value <0.05 ordered by enrichment values, calculated as the portion of genes under selection in the processes over the portion of genes significant in all GO-BPs. Points are color-filled by the adjusted p-value (p.adjust) and sized according to the gene numbers including in the corresponding GO-BP. (C) Gene lineage network plot shows the correlation among differentially expressed genes (DEGs) and GO-BP. Points are color-filled by the $\log_2$ fold change ($Log_2FC$) of genes compared between CHOW and Vehicle mice.

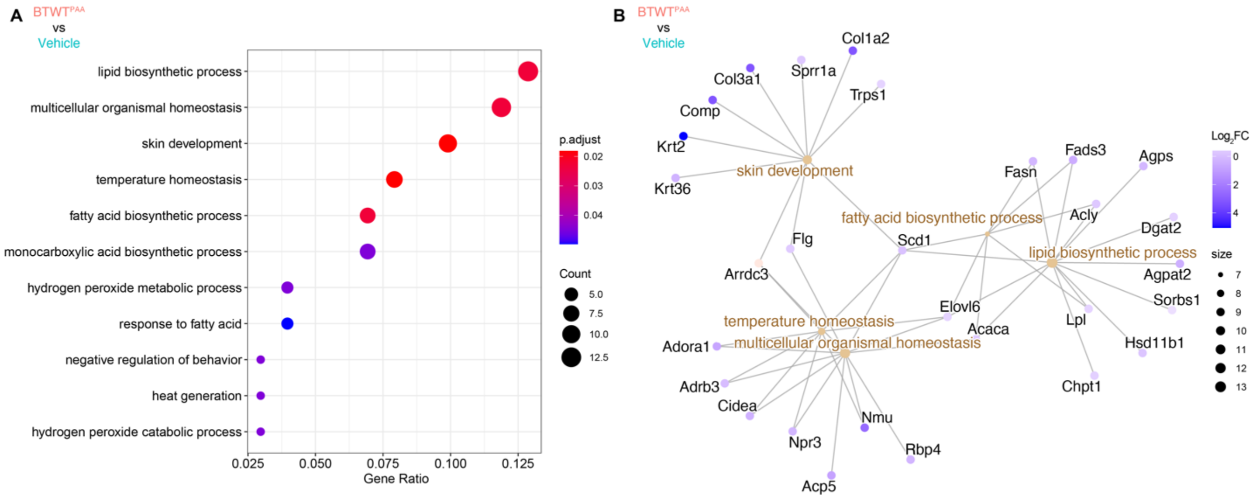


**Supplemental Figure 2.** Genes involved in lipid metabolism were enriched in the skin of mice that received sphingolipid-producing *B. thetaiotaomicron* ($BTWT^{PAA}$) compared to the mice fed solely lipid-free diet (Vehicle). (A) Biological processes in gene-ontology analysis (GO-BP) that are that are significant at the adjusted p-value (p.adjust), p <0.05, ordered by enrichment values, calculated as the portion of genes under selection in the processes over the portion of genes significant in all GO-BPs. Points are color-filled by p.adjust and sized according to the numbers of genes included in the corresponding GO-BP. (B) Gene lineage network plot shows the correlation among DEGs and GO-BP. Points are color-filled by the $\log_2$ fold change ($Log_2FC$) of genes compared between BTWT and Vehicle.

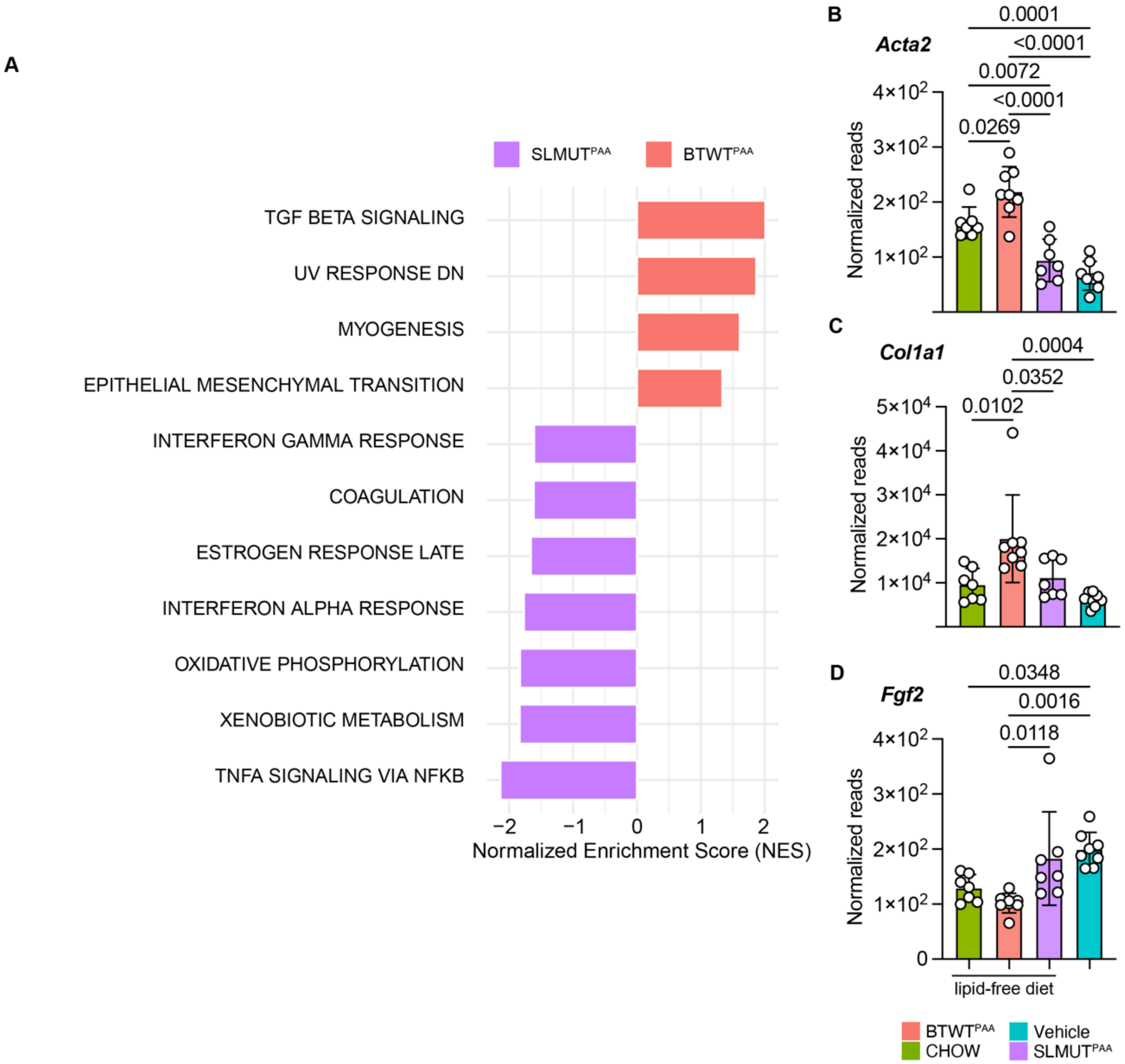


**Supplemental Figure 3.** Genes involved in TGF-β signaling and epithelial mesenchymal transition (EMT) were enriched in the skin of mice exposed to sphingolipid-competent versus sphingolipid-deficient *B. thetaiotaomicron.* Mice received either chow (CHOW) or a lipid-free diet; and mice fed with lipid-free diet were orally gavaged with either Vehicle, BTWT$^{PAA}$, or SLMUT$^{PAA}$. (n = 8 in either BTWT$^{PAA}$ or Vehicle; n = 7 in either CHOW or SLMUT$^{PAA}$) (A) Enriched Hallmark gene sets in either BTWT$^{PAA}$ or SLMUT$^{PAA}$. (B-D) Comparison of gene expression levels of (B) *Acta2* and (C) *Col1a1* in the TGF-β signaling pathway and (D) *Fgf2* in the antagonistic fibroblast growth factor 2 program.

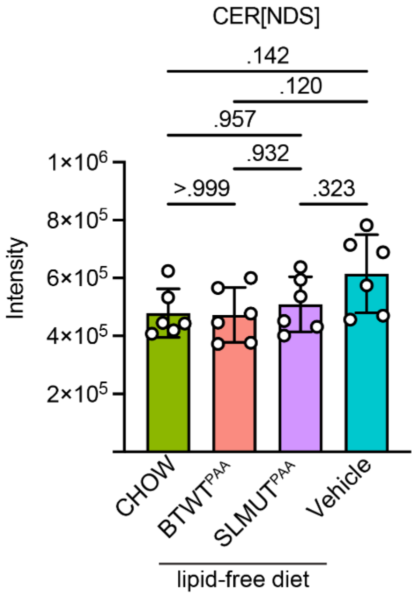


**Supplemental Figure 4.** No difference of the level of total CER[NDS] was found between groups. Mice received either chow (CHOW) or a lipid-free diet; and mice fed with lipid-free diet were orally gavaged with either a vehicle control (Vehicle), $BTWT^{PAA}$, or $SLMUT^{PAA}$. Total CER[NDS] levels in each group were calculated by summing up the sphingolipid species identified in this class. (n = 8 for both $BTWT^{PAA}$ and Vehicle; n = 7 for both CHOW and $SLMUT^{PAA}$)

**Supplemental Table 1.** SRM analysis

| Ceramide Species | Precursor ions (Q1) | | Product ion (Q3) | Collision energy (eV) |
|---|---|---|---|---|
| | **[M–H2O+H]+** | **$[M+H]^+$** | | |
| **CER[NDS]** | | | | |
| NDS (C16:0) | | 540.5 | 284.3 | 30 |
| NDS (C20:0) | | 596.6 | 284.3 | 30 |
| NDS (C24:0) | | 652.6 | 284.3 | 30 |
| NDS (C26:0) | | 680.7 | 284.3 | 30 |
| NDS (C28:0) | | 708.7 | 284.3 | 30 |
| NDS (C30:0) | | 736.7 | 284.3 | 30 |
| NDS (C32:0) | | 764.8 | 284.3 | 40 |
| NDS (C34:0) | | 792.8 | 284.3 | 40 |
| NDS (C36:0) | | 820.8 | 284.3 | 40 |
| **CER[NS]** | | | | |
| NS (C14:0) | 492.4 | | 264.3 | 30 |
| NS (C18:0) | 548.5 | | 264.3 | 30 |
| NS (C22:0) | 604.6 | | 264.3 | 30 |
| NS (C26:0) | 660.7 | | 264.3 | 30 |
| NS (C32:0) | 744.8 | | 264.3 | 40 |
| NS (C36:0) | 800.8 | | 264.3 | 40 |
| **CER[EOS]** | | | | |
| EOS (C26:0) | 938.9 | 956.9 | 264.3 | 30 |
| EOS (C30:0) | 995.0 | 1013.0 | 264.3 | 40 |
| EOS (C34:0) | 1051.1 | 1069.1 | 264.3 | 40 |
| EOS (C26:1) | 936.9 | 954.9 | 264.3 | 30 |
| EOS (C30:1) | 993.0 | 1011.0 | 264.3 | 40 |
| EOS (C34:1) | 1041.1 | 1067.1 | 264.3 | 40 |
| **CER[OS]** | | | | |
| OS/P-OS (C30:0) | 732.7 | 750.7 | 264.3 | 30 |
| OS/P-OS (C32:0) | 760.8 | 778.8 | 264.3 | 40 |
| OS/P-OS (C34:0) | 788.8 | 806.8 | 264.3 | 40 |
